# Brain-Penetrating Peptide and Antibody Radioligands for Proof-of-Concept PET Imaging of Fibrin in Alzheimer’s Disease

**DOI:** 10.1101/2025.06.02.657356

**Authors:** Dag Sehlin, Ximena Aguilera, Marta Cortés-Canteli, Stina Syvänen, Sara Lopes van den Broek

## Abstract

**Background:** Alzheimer’s disease (AD) is increasingly recognized as a multifactorial disorder with vascular contributions, including a pro-coagulant state marked by fibrin deposition in the brain. Fibrin accumulation may exacerbate cerebral hypoperfusion, leading to neurodegeneration. Identifying patients with this pathology could enable targeted anticoagulant therapy. However, current imaging tools lack the specificity and sensitivity to detect fibrin in the brain. This study aimed to develop and evaluate brain-penetrating peptide- and antibody-based PET radioligands targeting fibrin to enable individualized treatment strategies in AD.

**Results:** A fibrin-binding peptide (FBP) was conjugated to the antibody fragment scFv8D3, which targets the transferrin receptor (TfR), to facilitate transcytosis across the blood-brain barrier. FBP-scFv8D3 bound TfR and with modest affinity to fibrin, though with limited selectivity over fibrinogen. In vivo studies in Tg-ArcSwe mice, that exhibit fibrin along with brain amyloid-β pathology, and wild-type mice showed that [^125^I]FBP-scFv8D3 retained brain-penetrating properties but did not demonstrate significant fibrin-specific retention. In contrast, the monoclonal antibody 1101 and its bispecific, brain penetrant variant 1101-scFv8D3 exhibited high fibrin selectivity and TfR binding. Both antibodies showed a trend towards higher brain retention in Tg-ArcSwe mice and [^125^I]1101-scFv8D3 showed a higher brain-to-blood ratio compared to [^124^I]1101. PET imaging with [^124^I]1101 and [^124^I]1101-scFv8D3 revealed low brain uptake but ex vivo autoradiography suggested specific cortical retention in Tg-ArcSwe mice.

**Conclusion:** This study demonstrates the feasibility of using bispecific antibody-based PET radioligands to target fibrin in the AD brain. While the FBP-scFv8D3 conjugate showed limited specificity, the bispecific antibody 1101-scFv8D3 exhibited promising brain penetration and fibrin selectivity. These findings support further development of antibody-based imaging tools toward the goal to stratify AD patients who may benefit from anticoagulant therapy.

## Introduction

Alzheimer’s disease (AD) neuropathology is characterized by amyloid plaques, tau tangles, and neuro inflammation, which leads to neurodegeneration and cerebral atrophy.^1,2^ In recent years, AD has also been linked with vascular risk factors with accumulating evidence showing a decreased cerebral blood flow in AD patients. More specifically, the level of cerebral hypoperfusion in the AD brain has been correlated to the degree of dementia. This may be partially due to a significant pro-coagulant state present in AD, driven by the formation and deposition of fibrin, the main protein of blood clots.^3–5^ Fibrin is found both in the vasculature, where it builds up from the blood, and inside the brain, in the parenchyma. The amount of fibrin pathology appears to be highly variable within AD patients, which can be explained by the heterogeneity of the AD population. The pro-coagulant state is increasingly recognized as a factor in AD progression, and anticoagulant therapies aimed at reducing fibrin(ogen) levels may offer a promising strategy to normalize cerebral blood flow.^6^ However, since this pro-coagulant state is only present in a subset of patients, individualized treatment approaches are essential. Taking into consideration that anticoagulants are already widely available, their repurposing for AD treatment is both feasible and cost-effective, particularly in comparison to newly approved AD immunotherapies.^7,8^ To identify the patients that can benefit from anticoagulant treatment, novel sensitive and specific diagnostic tools must be developed.^9^

Positron emission tomography (PET) is a non-invasive nuclear molecular imaging technique that enables sensitive quantification and visualization of disease targets.^10–14^ Currently, most PET radioligands are small molecules. To cross the blood-brain barrier (BBB), radioligands must be sufficiently lipophilic, which can result in off-target distribution within brain tissue. Moreover, small molecule radioligands often exhibit limited target specificity and sensitivity, reducing the reliability and accuracy of PET imaging.

Peptide- and antibody-based PET ligands offer a promising alternative due to their high target specificity, selectivity, and low non-displaceable binding.^15–18^ However, their relatively large size prevents them from efficiently crossing the BBB. This limitation underscores the need for innovative strategies to enhance brain uptake. One of the most widely explored approaches is receptor-mediated transcytosis, which employs bispecific constructs that bind both the intrabrain target molecule and a receptor expressed on the BBB, such as the transferrin receptor (TfR), which facilitates transcytosis of the ligand across the BBB and into the brain.^19–22^

In this study, we aim to develop brain-penetrating fibrin-targeting PET radioligands capable of detecting the pro-coagulant state in AD. More specifically, we developed a bispecific fibrin-binding peptide as well as a bispecific antibody ligand and evaluated their potential to identify fibrin in the brain, with a future aim to individualize treatment for AD patients that can benefit from anticoagulant therapy.

## Results

This study investigated two approaches for developing PET radioligands to image fibrin in the living brain: one based on a fibrin-binding peptide (FBP) and the other on a fibrin-binding antibody (1101). Both FBP and 1101 were conjugated to the mouse transferrin receptor (mTfR)-binding Fab fragment 8D3 for facilitated brain delivery.

### Conjugation of FBP to scFv8D3

A fibrin binding peptide (FBP), previously used to detect fibrin in heart tissue was synthesized with an NHS-ester at the peptide N-terminus (Figure S1).^23,24^ The NHS-ester was used as a handle for conjugation to lysine residues on scFv8D3. This method has been widely used as a bioconjugation strategy and allows the attachment of multiple probes to one antibody or antibody fragment.^25,26^ This method’s lack of selectivity imposes a risk of interference with the binding properties of the antibody (fragment) when conjugations are made within the binding region (CDR region).^27^ Therefore, scFv8D3 was designed with a lysine-rich FlagTag the C-terminus to steer the site of modification away from the binding regions.^28^ The conjugation efficiency was tested using different FBP-NHS ester:scFv8D3 ratios, using 15-50 equivalents (eq.) FBP-NHS excess.

### FBP-scFv8D3 binding to fibrin

Enzyme-linked immunosorbent assay (ELISA) was performed to confirm conjugation of FBP to scFv8D3 and assess the potential impact on the affinity of three FBP-scFv8D3 conjugates (15, 30 and 50 eq.) towards fibrin, fibrinogen and mTfR using a commercially available anti-fibrinogen antibody and scFv8D3 as controls (Figure 1). The anti-fibrinogen antibody showed equal affinity towards fibrin and fibrinogen, with no binding towards mTfR, whereas scFv8D3 bound to mTfR but not to fibrin or fibrinogen (Figure 1). All three FBP-scFv8D3 constructs bound fibrin, however, with substantially lower affinity compared to the control antibody. The FBP-scFv8D3 constructs exhibited binding to mTfR with affinities comparable to that of unmodified scFv8D3, indicating that FBP was successfully conjugated without compromising mTfR binding. FBP-scFv8D3 (50 eq.) showed a slightly higher binding towards fibrin than the other two versions, but also a somewhat reduced mTfR binding. This can be explained by the addition of more FBPs to the scFv8D3 moiety, which increases the fibrin binding but can also interfere with the scFv8D3 binding to mTfR. Notably, all FBP-scFv8D3 also showed some degree of binding to fibrinogen, which implies that conjugation to scFv8D3 may reduce the peptide’s affinity to fibrin and thereby the selectivity towards fibrin over fibrinogen.

**Figure 1.**
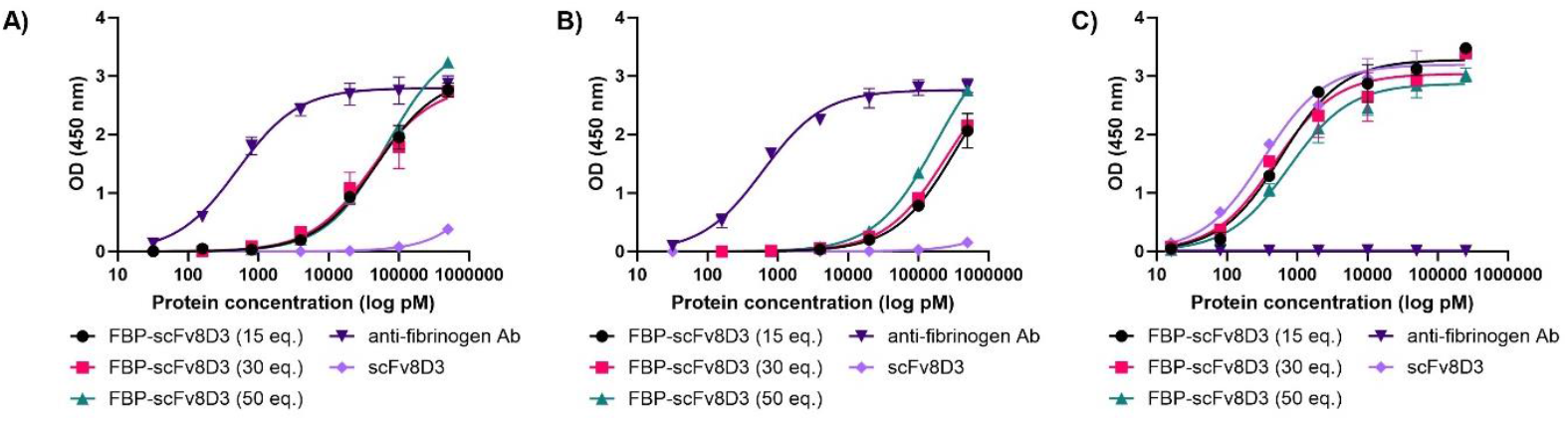
Indirect ELISA results showing FBP-scFv8D3 binding towards **A**) fibrin, **B**) fibrinogen and **C**) mTfR. The anti-fibrinogen antibody showed much higher binding towards both fibrin and fibrinogen compared to the FBP-scFv8D3 constructs and no binding towards mTfR, whereas scFv8D3 bound only to mTfR and not to fibrin or fibrinogen.

### Ex vivo evaluation FBP-scFv8D3

Brain uptake and biodistribution studies were performed with FBP-scFv8D3, conjugated at 25 eq. peptide excess. After iodine-125 (^125^I)-radiolabeling of FBP-scFv8D3, brain and blood concentrations were first evaluated in WT mice at 2 h after injection, using the non-conjugated scFv8D3 as control. Ex vivo analyses in WT animals showed similar brain uptake and brain-to-blood ratio for [^125^I]FBP-scFv8D3 and [^125^I]scFv8D3 at 2 h after injection, indicating that the addition of FBP moieties to the scFv8D3 did not significantly interfere with the brain penetrating capacities of scFv8D3 (Figure 2A-B). Analysis of biodistribution to peripheral organs indicated high concentrations of both ligands in spleen, consistent with previous studies on TfR binding ligands (Figure 2C). Next, [^125^I]FBP-scFv8D3 was evaluated in the Tg-ArcSwe mouse model of Aβ pathology, which has fibrin deposits associated both with parenchymal plaques and the brain the vasculature (Figure 2D). Measurements of [^125^I]FBP-scFv8D3 blood concentrations over 24 h showed no difference between Tg-ArcSwe and WT mice (Figure 2E). Brain concentrations of [^125^I]FBP-scFv8D3 measured 24 h post injection were overall low, with no difference between Tg-ArcSwe and WT mice (Figure 2F). Similarly, no difference was seen in the brain-to-blood ratio (Figure 2G), implying no significant binding to fibrin clots in the brain of Tg-ArcSwe mice.

**Figure 2.**
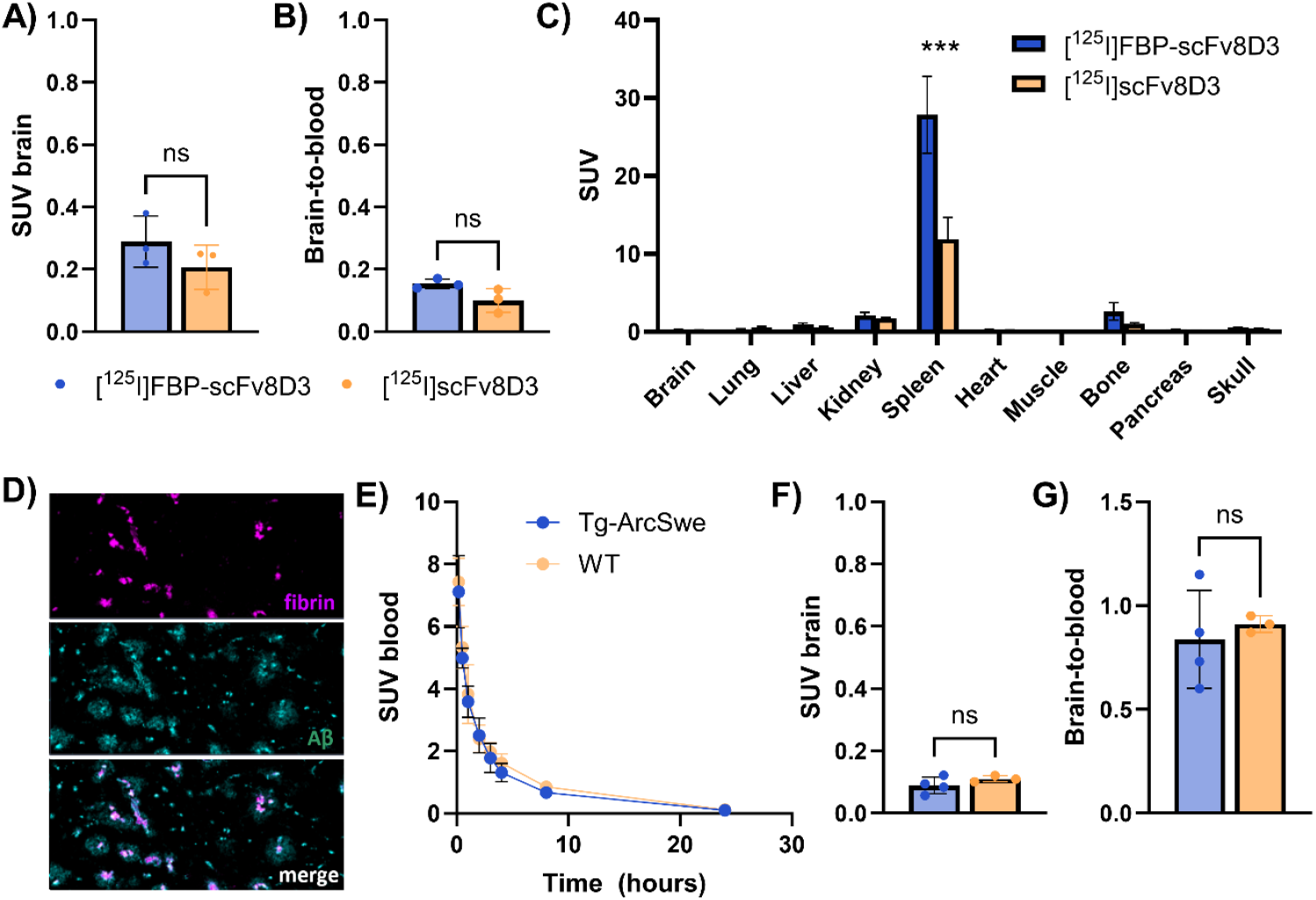
Ex vivo evaluation of [^125^I]FBP-scFv8D3. **A**) Brain uptake, expressed as standardized uptake value (SUV), of [^125^I]FBP-scFv8D3 and [^125^I] scFv8D3 in WT mice. **B**) Brain-to-blood ratio in WT mice 2 h after injection of [^125^I]FBP-scFv8D3 and [^125^I]scFv8D3. **C**) [^125^I]FBP-scFv8D3 and [^125^I]scFv8D3 biodistribution to peripheral organs at 2 h after injection in WT mice. **D**) Immunofluorescent staining of fibrin (magenta; antibody 1101) in combination with Aβ staining (cyan; LCO HS-84), confirming the presence of fibrin in Aβ plaques and brain vessels. **E**) Blood concentrations of [^125^I]FBP-scFv8D3 in Tg-ArcSwe and WT mice over 24 h. **F**) Brain uptake of [^125^I]FBP-scFv8D3 at 24 h after injection in Tg-ArcSwe and WT mice. **G**) Brain-to-blood ratio of [^125^I]FBP-scFv8D3 at 24 h after injection Tg-ArcSwe and WT mice.

### Fibrin selective antibodies 1101 and 1101-scFv8D3

While a relatively small peptide may be greatly affected by conjugation to scFv8D3, antibodies generally tolerate such fusions. Thus, the monoclonal fibrin-specific antibody 1101 was evaluated along with its bispecific variant 1101-scFv8D3, where scFv8D3 was attached to each of the light chains of antibody 1101, to enhance its brain uptake^29^. ELISA assays showed highly selective binding of antibody 1101 and 1101-scFv8D3 towards fibrin over fibrinogen, with similar affinity as the anti-fibrinogen antibody control (Figure 3A-B). 1101-scFv8D3 also bound to mTfR, while 1101 did not (Figure 3C).

**Figure 3.**
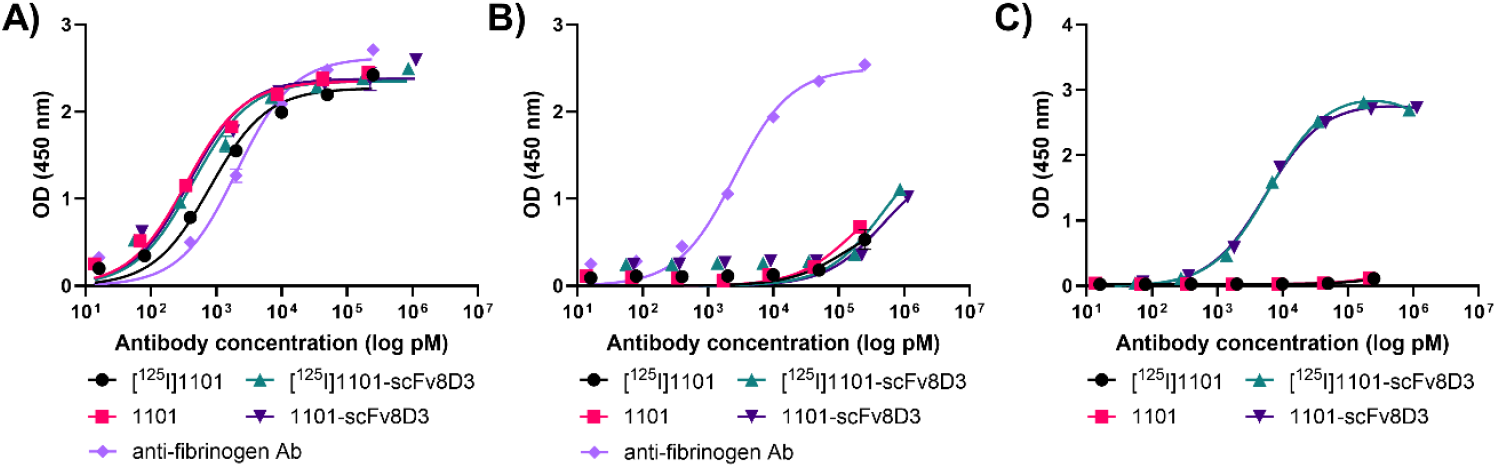
In vitro evaluation of 1101 and 1101-scFv8D3. 1101 and 1101-scFv8D3 were evaluated, before and after radiolabeling, with indirect ELISA for their binding towards **A**) fibrin; **B**) fibrinogen; and **C**) mTfR. A commercially available anti-fibrinogen antibody was used as control in **A** and **B**.

Antibodies 1101 and 1101-scFv8D3 were radiolabeled with iodine-125 and injected in WT mice. Both the brain uptake (Figure 4A) and the brain-to-blood ratio (Figure 4B) were significantly higher for the bispecific 1101-scFv8D3, indicating better brain-penetrating properties of the bispecific antibody. Biodistribution to peripheral organs (Figure 4C) showed higher uptake in the spleen of the bispecific antibody, again indicating TfR binding in the spleen. Next, [^125^I]1101 and [^125^I]1101-scFv8D3 were evaluated in the Tg-ArcSwe mouse model of Aβ pathology, which has fibrin deposits associated both with parenchymal plaques and the brain the vasculature (Figure 4D). In line with previous observations for TfR-binding bispecific antibodies, [^125^I]1101-scFv8D3 showed a rapid clearance from blood, significantly reducing its systemic exposure compared with [^125^I]1101 (Figure 4D). At 72 h post administration, similar brain retention was observed for both antibodies, with no significant differences between Tg-ArcSwe and WT mice, although both antibodies showed a trend towards higher retention in Tg-ArcSwe mice (Figure 4E). Due to its fast blood clearance, [^125^I]1101-scFv8D3 displayed a significantly higher brain-to-blood ratio compared with [^125^I]1101, and also a significant difference between Tg-ArcSwe and WT mice (Figure 4F). The brain-to-blood ratio is especially relevant for PET imaging, where high imaging contrast is dependent on the specific binding to the target (brain) vs low background (blood).

**Figure 4.**
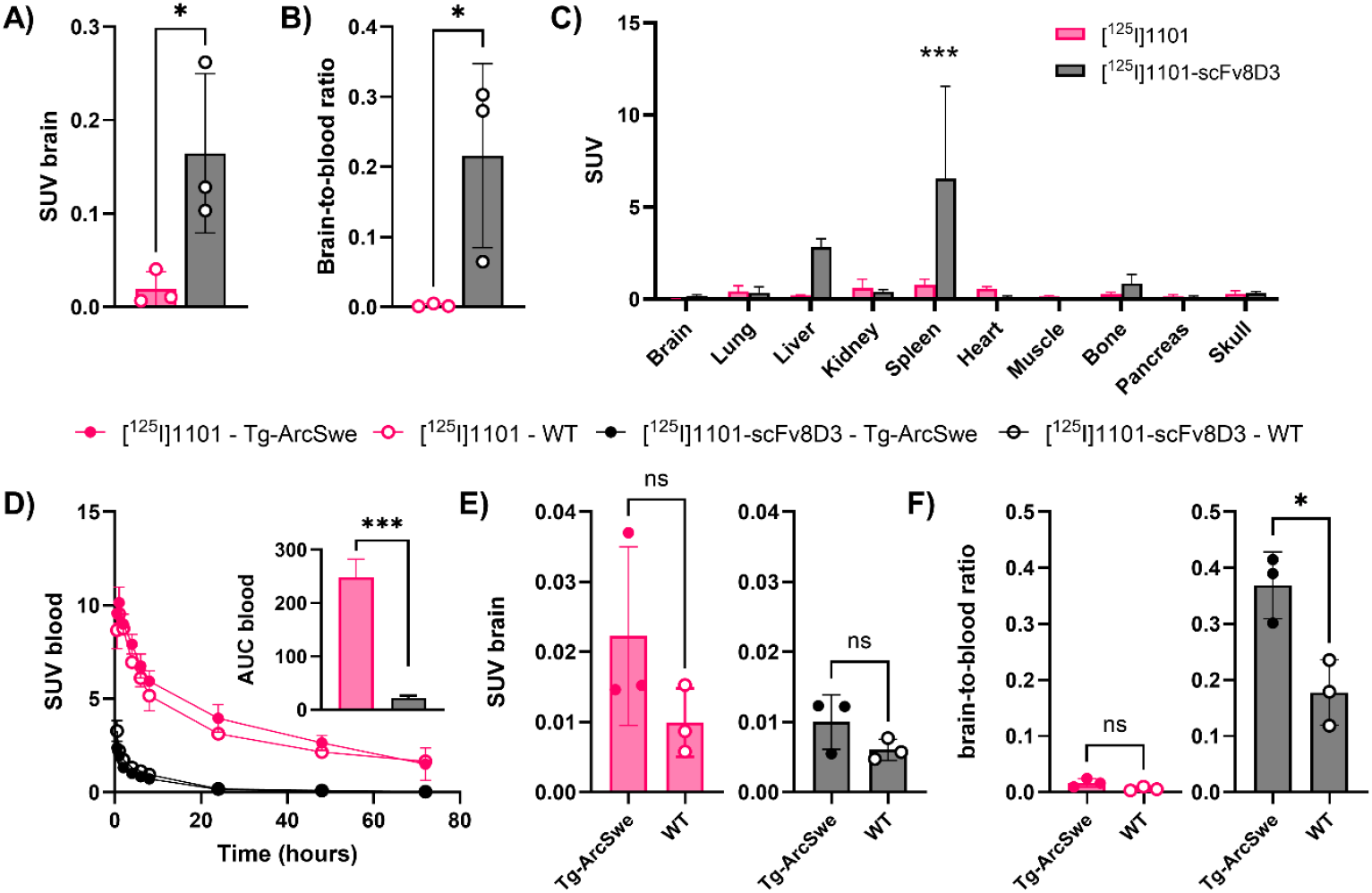
Ex vivo evaluation of [^125^I]1101 and [^125^I]1101-scFv8D3. **A**) Brain uptake in WT mice 2 h after injection of [^125^I]1101 and [^125^I]1101-scFv8D3. **B**) Blood-to-brain ratio in WT mice after 2 h after injection of [^125^I]1101 and [^125^I]1101-scFv8D3. **C**) Biodistribution of [^125^I]1101 and [^125^I]1101-scFv8D3 in WT mice at 2 h after injection. **E**) Blood concentration of [^125^I]1101 and [^125^I]1101-scFv8D3 in Tg-ArcSwe and WT mice over 72 h. Inset: Area under the curve for [^125^I]1101 and [^125^I]1101-scFv8D3 blood concentration between 0 and 72 h. **F**) Brain concentrations and **G**) brain-to-blood ratio of [^125^I]1101 and [^125^I]1101-scFv8D3, 72 h after injection in Tg-ArcSwe and WT mice.

### PET imaging with 1101 and 1101-scFv8D3

Antibodies 1101 and 1101-scFv8D3 were radiolabeled with iodine-124 (^124^I) and administered to Tg-ArcSwe and WT mice for immunoPET imaging. PET images acquired 72 ha post administration revealed overall low brain concentrations of both antibodies, relative to surrounding tissues, with no clear differences between Tg-ArcSwe and WT mice. Ex vivo autoradiography images, however, suggested a specific retention of both antibodies in the brains of Tg-ArcSwe mice, most notably in the cortex (Figure 5, Figure S2). Quantification of PET data confirmed the visual impression from PET images, with equal average brain concentrations in Tg-ArcSwe and WT mice (Figure 5B). In line with the autoradiography, ex vivo quantification of radioactivity in post mortem tissue revealed elevated concentrations and brain-to-blood ratios of both antibodies in Tg-ArcSwe mice compared to WT. However, due to variation and the low number of animals, differences were not statistically significant (Figure 5C).

**Figure 5.**
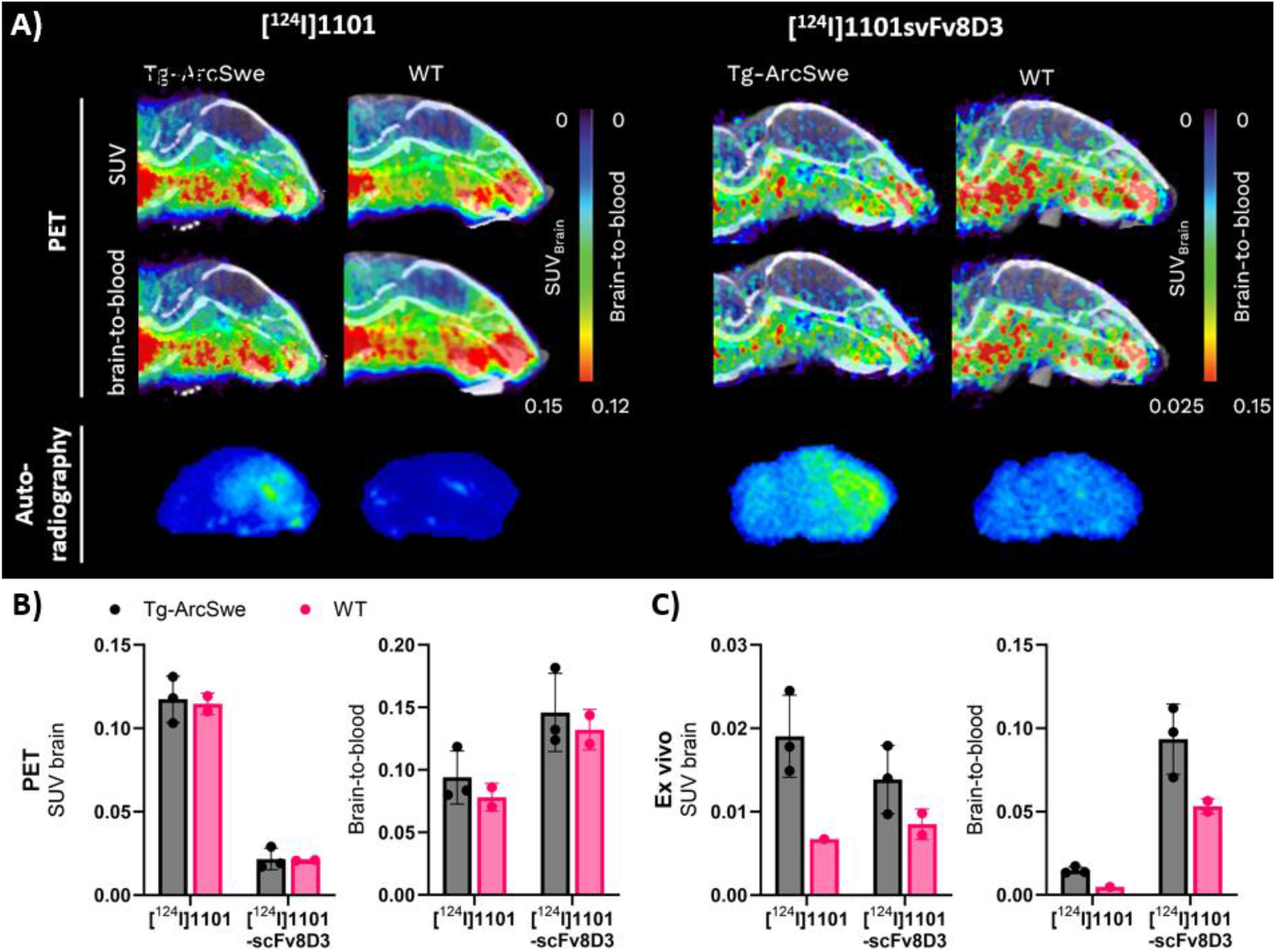
ImmunoPET with [^124^I]1101 and [^124^I]1101-scFv8D3. **A)** Representative sagittal PET images of Tg-ArcSwe and WT mice acquired 72 h post injection of [^124^I]1101 or [^124^I]1101-scFv8D3, expressed as standardized uptake value (SUV, upper panel) or brain to blood ratio (middle panel). Note the different scales for the two antibody ligands. Ex vivo autoradiography (lower panel) from saline perfused, PET scanned mice. **B)** Quantification of PET images from **A**), expressed as SUV and brain-to-blood concentration ratio. **C**) Ex vivo quantification of [^124^I]1101 and [^124^I]1101-scFv8D3 concentration in brain tissue from PET scanned mice in **A**), expressed as SUV and brain-to-blood concentration ratio.

## Discussion

This study presents a significant advancement in the development of molecular imaging tools for Alzheimer’s disease (AD), particularly targeting the vascular component of the disease through fibrin imaging. The rationale stems from increasing evidence that a subset of AD patients exhibit a pro-coagulant state characterized by fibrin deposition, which contributes to neurovascular dysfunction, impaired cerebral perfusion, and potentially accelerates neurodegeneration. Detecting this pathology non-invasively could enable stratification of patients for anticoagulant therapy, a promising and cost-effective intervention given the widespread availability of such drugs.

The peptide-based radioligand FBP-scFv8D3 was designed to combine fibrin specificity with brain-penetrating capabilities via TfR-mediated transcytosis. While the construct retained its ability to cross the BBB, as evidenced by comparable brain uptake to the parent scFv8D3, its binding affinity to fibrin was modest and lacked sufficient selectivity over fibrinogen. It is likely that conjugation of the FBP with the relatively larger scFv8D3 negatively affected its fibrin binding. This limitation is critical, as high off-target binding in the periphery could compromise imaging specificity. Moreover, in vivo studies did not show significant fibrin-specific retention in Tg-ArcSwe mice. Taken together, the affinity and selectivity of the FBP-scFv8D3 conjugate was insufficient for effective in vivo imaging.

In contrast, the monoclonal antibody 1101 and its bispecific variant 1101-scFv8D3 demonstrated high affinity and selective binding for fibrin over fibrinogen. The bispecific format enabled efficient BBB transcytosis, yielding substantial increase in brain uptake compared to the monospecific 1101 antibody, without compromising target binding. The brain uptake at 2 h post-injection was somewhat lower for 1101-scFv8D3 compared to the FBP-scFv8D3 peptide conjugate, which could reflect the larger size of the antibody and correspondingly slower kinetics. Notably, initial ex vivo studies showed a trend for both antibodies towards higher brain retention in Tg-ArcSwe over WT mice, indicating potential for specific fibrin imaging. The bispecific [^125^I]1101-scFv8D3 also exhibited an elevated brain-to-blood ratio, which was significantly higher in Tg-ArcSwe compared to WT mice. Although PET imaging showed low absolute brain uptake, ex vivo autoradiography revealed cortical retention in transgenic mice, primarily in the cortical region, which is the primary site for development of protein pathology in this model, supporting the hypothesis of target engagement.

These findings underscore the importance of molecular format in radioligand design. While peptides offer advantages in terms of size and synthetic flexibility, their binding characteristics may be more susceptible to modification upon conjugation. Antibodies, particularly when engineered as bispecific constructs, provide a more robust platform for achieving both target specificity and brain delivery. The use of TfR-mediated transcytosis is particularly promising, as it allows for the delivery of large biologics across the BBB, a major hurdle in CNS drug and imaging agent development. It should be noted, however, that fibrin is likely present on both the luminal and abluminal sides of the BBB. Therefore, a conventional IgG without TfR-mediated transcytosis could still be sufficient, which may explain the relatively small differences observed in brain retention between [^125^I]1101 and [^125^I]1101-scFv8D3. Nonetheless, the faster systemic clearance of the bispecific variant, [^125^I]1101-scFv8D3, is advantageous from a PET imaging perspective, as it contributes to more rapid background clearance and improved contrast between specific and non-specific signals. However, several limitations must be addressed in future studies. First, the sample sizes in animal experiments were relatively small, limiting statistical power. Increasing the number of animals and including additional time points could provide more definitive evidence of target engagement and pharmacokinetics. Here, it was not possible to increase the number of included animals since the antibody proved to be very difficult to express. Thus, other expression systems, or other antibody formats, should be investigated for obtaining larger quantities of the antibodies. Second, the radiolabeling strategy and choice of isotope may influence imaging sensitivity and resolution. Optimization of radiochemistry, including site-specific labeling and use of isotopes with more favorable decay properties, could enhance imaging performance.

Additionally, while the Tg-ArcSwe model recapitulates key aspects of AD pathology, including variable degrees of fibrin deposition, it remains primarily a model of Aβ pathology, and thus a simplified representation of the human disease. Validation in other models and, ultimately, in human tissue or clinical studies will be essential to confirm the translational potential of this radioligand.

## Conclusion

In conclusion, this study provides a compelling proof-of-concept for the use of bispecific antibody-based PET radioligands to detect fibrin in the AD brain. The results support further development and optimization of such tools for patient stratification and monitoring of anticoagulant therapy. By enabling personalized treatment approaches, this strategy could significantly improve outcomes for a subset of AD patients and represents a promising direction in the evolving landscape of precision medicine in neurodegenerative diseases.

## Materials and Methods

### Conjugation of FBP to scFv8D3

Single chain fragment variable scFv8D3-FlagTag was produced according to previously published procedures.^28^ FBP functionalized with an NHS ester at the C-terminal was produced on special request by ThermoFischer. FBP was stored in DMSO and freshly diluted in PBS before the conjugation reaction. Sodium carbonate buffer (1M, pH 8.0) was added to an Eppendorf tube containing scFv8D3-FlagTag (1-1.20 mg/mL in PBS, 20-100 uL) to give a final concentration of 30 mM. FBP dissolved in PBS containing. 10% DMSO was added to the mixture using 5-100 equiv. peptide-scFv8D3 ratio. The mixture was incubated at room temperature for 2 hours at 600 RMP. The solution was purified to remove unreacted FBP using Zeba spin desalting columns (7K MWCO, 0.5 mL, 89882, Thermofisher) and eluted in PBS (pH 7.4). Final protein concentration was measured with spectrophotometry (DeNovix spectrophotometer DS-11).

### Expression of antibodies 1101 and 1101-scFv8D3

Antibody 1101 was designed as a mouse IgG2c backbone with the variable domains from a previously described fibrin specific antibody. ^30^ Its bispecific variant 1101-scFv8D3 was designed with scFv8D3 fused to the C-terminal ends of the 1101 light chains. Both antibodies were expressed in Expi293 cells and purified using protein A chromatography, according to a previously described protocol.^29^

### SDS-PAGE

SDS-PAGE analyses were conducted to confirm the size and integrity of the antibodies and FBP-scFv8D3 conjugates. The protein samples were mixed with Bolt LDS sample buffer (ThemoFischer, 13276499), heated for 2 min at 95°C, and loaded onto 4-12% Bolt Bis-Tris Plus Gels (ThermoFischer) and run for 22 min at 200 V in MES buffer (ThermoFischer, 13266499). PageRuler™ Plus Prestained Protein was added as a molecular weight standard (ThermoFischer, 11832124). Afterwards, the gel was removed and washed in dH2O and stained with InstantBlue (Abcam, ab119211). Images of the gels were taken with a ChemiDoc XRS+ Gel Imaging System (BioRad) and analyzed with ImageLab (BioRad).

### ELISA

**Fibrin**. Fibrin ELISA assay methodology was adapted from Hanaoka et al.^32^ The 96-well half-area plates (Corning Inc., New York, NY) were coated overnight at 4°C with fibrinogen (50 µL/well, 10 µg/mL diluted in PBS, Sigma-Aldrich F3879). A mixture of 0.05 U/mL thrombin (Sigma-Aldrich 605195), 7mM L-cysteine and 1 mM CaCl_2_ with a total volume of 50 µL/mL was added and the plates were incubated for 2h at 37°C. **Fibrinogen**. The 96-well half-area plates (Corning Inc., New York, NY) were coated overnight at 4°C with fibrinogen (50 µL/well, 10 µg/mL diluted in PBS, Sigma-Aldrich F3879). **Mouse transferrin receptor**. The 96-well half-area plates (Corning Inc., New York, NY) were coated overnight at 4°C with mTfR (50 µL/well, 5 µg/mL diluted in PBS). **Blocking**. The wells were blocked with 1% BSA in PBS for 1h. The proteins were diluted in incubation buffer (PBS containing 0.1% BSA, 0.05% Tween-20 and 0.15% Kathon) **Primary antibody**. The plates were washed with wash buffer (phosphate buffer with 0.1% Tween-20 and 0.15% Kathon, pH 7.5). 1101, 1101-scFv8D3, FBP-scFv8D3, their radiolabeled equivalent, and control antibodies were serial diluted in incubation buffer (PBS with 0.1% BSA, 0.05% Tween-20 and 0.15% Kathon) from 250 nM (for fibrin and fibrinogen coated plates) or 50 nM (for mTfR and mIgG coated plates) to 3.2 pM and incubated for 2 hours. **Secondary antibody**. The plates were washed with wash buffer and detected with horseradish peroxidase (HRP)-coupled goat anti-mouse IgG-F(ab’)2 (1:2000, Jackson ImmunoResearch Laboratories, West Grove, PA, #115-035-006) or rabbit anti-goat IgG (1:4000, Thermofischer, #31402). **Plate development**. Signals were developed with K blue aqueous TMB substrate (Neogen Corp., Lexington, KY), quenched with 1M H_2_SO_4_ and read with a spectrophotometer at 450 nm. The data was analyzed using GraphPad Prism.

### Animals

The Tg-ArcSwe mouse model harbors the Arctic (E693G) and Swedish (KM670/671NL) APP mutations and is maintained on a C57BL/6 background. Tg-ArcSwe mice show elevated levels of soluble Aβ protofibrils at a young age and abundant and rapidly developing plaque pathology starting at around 6 months of age.^33,34^ Both males and females were used and WT littermates were used as control animals. The animals were housed in rooms with controlled temperature and humidity in an approved facility at Uppsala University with *ad libitum* access to food and water. All procedures described in this paper were approved by the Uppsala County Animal Ethics board (5.8.18-20401/2020), following the rules and regulations of the Swedish Animal Welfare Agency and in compliance with the European Communities Council Directive of 22 September 2010 (2010/63/EU).

### Radiochemistry

FBP-scFv8D3 and the two antibodies 1101 and 1101-scFv8D3 were radiolabeled by direct radioiodination of iodine-125 using the Chloramine-T method.^35^ FBP-scFv8D3 (70 µg, 58.3 µL, 2500 pmol), 1101 (70 µg, 25.5 µL, 467 pmol) or 1101-scFv8D3 (100 µg, 102 µL, 476 pmol) was mixed with 13.7-15.4 MBq [^125^I]-NaI (Perkin-Elmer Inc Waltham, MA, USA) and 10 µg Chloramine-T (Sigma Aldrich, Stockholm, Sweden) and the reaction mixture was diluted in PBS to a final volume of 260 µL. The reaction was incubated for 90 s at room temperature and subsequently quenched with 20 µg sodium metabisulfite (Sigma Aldrich, Stockholm, Sweden). The product was diluted with PBS to 500 µL and unreacted iodine, Chloramine-T and sodium metabisulfite were removed with a NAP-5 size exclusion column (GE Healthcaer, Uppsala, Sweden) with a total elution volume of 1 mL PBS. The ^125^I-iodination resulted in molar activities of 5.3 MBq/nmol, 24.4 MBq/nmol, and 13.7 MBq/nmol for FBP-scFv8D3, 1101, and 1101-scFv8D3, respectively.

For PET imaging, 1101 and 1101-scFv8D3 were labeled with iodine-124. 120 µl [^125^I]-NaI stock solution was preincubated 15 min with 12 µl cold NaI (100 µM), then neutralized with 24 µl of 0.5% acetic acid and 17 µl 10xPBS before addition of 145 µg of 1101 or 160 µg of 1101-scFv8D3. Each reaction was started by the addition of Chloramine T to a final concentration of 75 µg/ml and stopped after 2 min with Na-metabisulphite at 150 µg/ml. Labeled antibodies were purified with NAP-5 columns as above. The ^124^I-iodination resulted in molar activities of 36.4 MBq/nmol for 1101 and 44.1 MBq/nmol for 1101-scFv8D3.

### Ex vivo biodistribution

Mice (3-24 months; n=28) were injected intravenously (i.v.) in the tail vein with either 1.0 MBq ± 0.15 [^125^I]FBPscFv8D3; 0.42 MBq ± 0.036 [^125^I]scFv8D3; 1.0 MBq ± 0.18 [^125^I]1101; or 0.56 MBq ± 0.11 [^125^I]1101-scFv8D3. Blood samples (8 µL) were obtained from the tail vein of selected individuals at 0.5, 1, 2, 5 4, 6, 8, 24, 48h to investigate the radioactivity concentration in blood over time. After 2, 24 or 72h, mice were anaesthetized with isoflurane and a terminal blood sample was taken from the heart, followed by transcardial perfusion with 40 mL of 0.9% NaCl for 2.5 to clear the brain and organs from blood. Thereafter, lung, liver, kidney, whole heart, pancreas, spleen, femoral muscle, femoral bone, skull bone and submandibular glands were isolated to evaluate the biodistribution of the radiolabeled compounds. The brain was divided into left and right hemispheres, and the left hemisphere was further divided into cerebrum and cerebellum. The brain samples were immediately frozen on dry ice. The radioactivity of the samples was measured with a gamma-counter (2480 Wizard™, Wallac Oy PerkinElmer, Turku, Finland). Protein concentrations were expressed as percent of injected dose per gram tissue (%ID/g) or as %ID/g corrected for body weight, referred to as standardized uptake value (SUV).

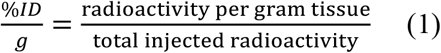

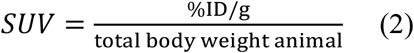

The brain-to-blood ratio was calculated by dividing the radioactivity in the brain per gram brain and the radioactivity in the blood per gram blood.

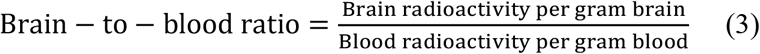

### Immunofluorescence

Immunofluorescence was performed to visualize Aβ pathology and localize the administered antibody in sagittal cryosections of mouse brain tissue. Tissue sections (20 µm thick) were initially fixed in ice-cold methanol for 10 min and subsequently rinsed twice in phosphate-buffered saline (PBS) for 5 min each. To block unspecific binding and permeabilize the tissue, sections were incubated for 2 h in a blocking solution containing 5% normal goat serum (NGS) in PBS. After blocking, sections were washed three times for 5 min in PBS. Antibody staining was performed by incubating the sections with goat anti-mouse Alexa Fluor 647-conjugated antibody (A21235, Invitrogen) diluted 1:200 in PBS 0.05% Tween-20 for 2 h at room temperature. Following this incubation, slides were washed twice for 5 min in PBS. To label Aβ fibrils, sections were then incubated with the luminescent conjugated oligothiophene (LCO) HS-84, diluted to a final concentration of 50 nM in PBS0.05% Tween-20, for 15 min at room temperature under gentle shaking. Excess dye was removed by washing the slides three times for 5 min in PBS. Sections were mounted using a mounting medium containing DAPI and imaged using a Zeiss Observer Z.1 microscope with ZEN 3.7 software (Carl Zeiss Microimaging GmbH, Jena, Germany).

### PET imaging

Tg-ArcSwe (n=3 per antibody) and WT (n=2 per antibody) mice were administered 5.2 ± 0.73 kBq/g and 6.8 ± 0.84 kBq/g body weight of [^124^I]1101 and [^124^I]1101-scFv8D3, respectively. At 72 h post-injection, a 120 min PET scan (Mediso NanoPET/MR, Mediso Medical Imaging System, Hungary) was acquired, followed by a 5-minute CT scan (Mediso NanoSPECT/CT, Mediso Medical Imaging System). Anaesthesia was maintained throughout the scanning procedure using 3.5–4.0% sevoflurane in a 0.5 L/min flow of 50% oxygen and 50% medical air. PET data were reconstructed on a 160 × 160 × 128 grid with voxel dimensions of 0.5 × 0.5 × 0.6 mm^3^ using three-dimensional ordered-subsets expectation maximization (20 iterations). CT data were reconstructed using filtered back-projection. Processing of the PET and CT images was performed with Amide, version 1.0.4.^36^ The CT scans were manually aligned with a T2-weighted, MRI-based mouse brain atlas.^37^ The PET images were subsequently aligned with the CT image, and a region of interest representing the cerebrum (brain) was outlined.

After the CT scan, a terminal blood sample was obtained from the heart before the mice were perfused, and the brain and peripheral organs were collected using the same procedure as described above. Radioactivity in the tissues was quantified via γ-counting, and radioactivity concentrations were converted to SUV. Sagittal brain sections (20 µm) were prepared from the right hemispheres and exposed to a phosphor imaging plate (MS, MultiSensitive, PerkinElmer) for 10 days. Plates were scanned in a Typhoon phosphor imager (Cytiva).

### Statistical analyses

Statistical analyses were performed in GraphPad Prism 10.2.2 (GraphPad Software, Inc., San Diego, CA). The results are reported as mean ± standard deviation and statistical assessment was conducted by unpaired t-test or a two-way ANOVA with multiple comparisons test.

## Declarations

### Ethics approval and consent to participate

Animal experiments were conducted under ethical permit number 5.8.18-20401/2020, issued by the Uppsala County Animal Ethics board.

### Consent for publication

*Not applicable*

### Availability of data and material

*The datasets used and/or analysed during the current study are available from the corresponding author on reasonable request*.

### Authors’ contributions

DS, MCC, SS and SLvdB designed the study. XA expressed the antibodies. DS and SLvdB performed radiolabeling. DS, SS and SLvdB performed ex vivo and PET imaging studies and analyzed the data. DS and SLvdB wrote the paper with input from all other authors.

## Acknowledgements

The molecular imaging work in this study was performed at the Preclinical PET-MRI Platform, a research infrastructure at Uppsala University, Sweden.

## Supplementary information

**Figure S1.**
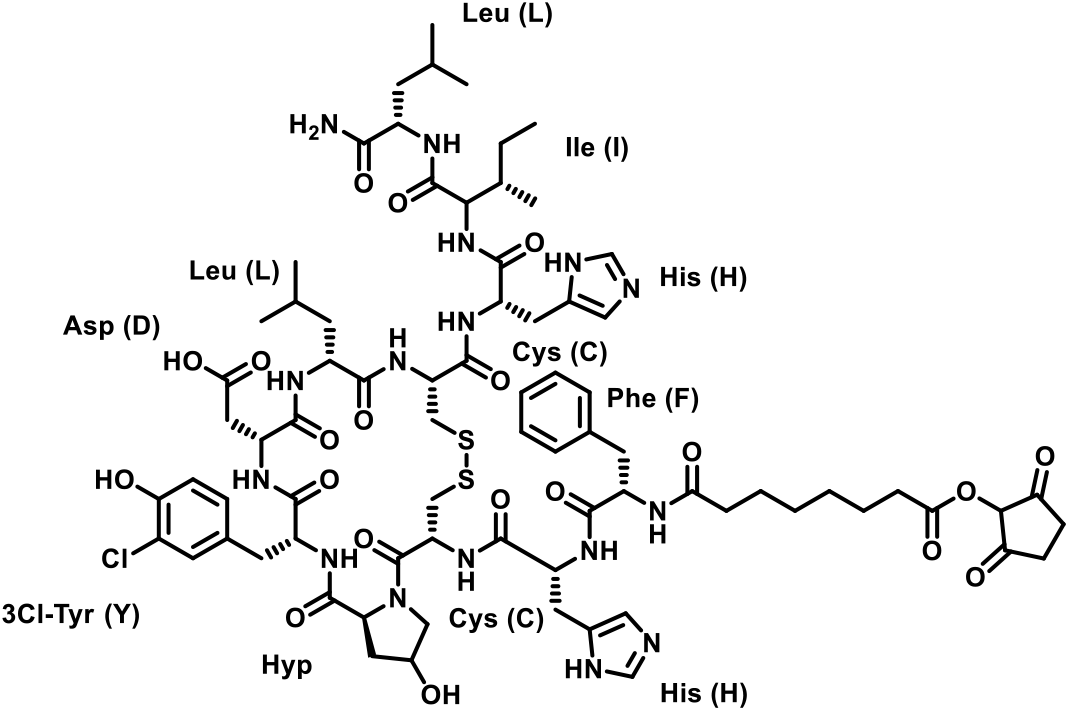
Structure of FBP.

**Figure S2.**
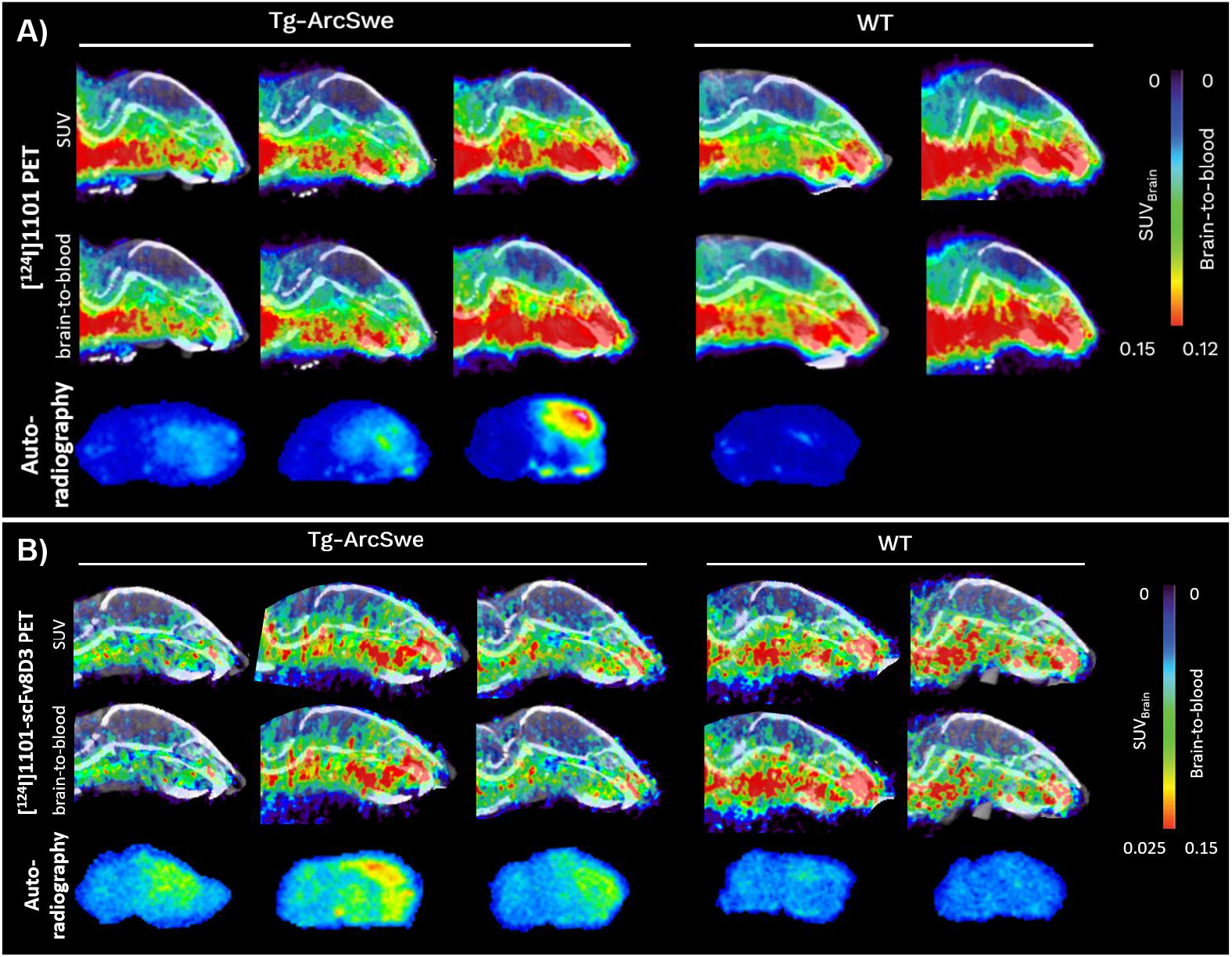
ImmunoPET with [^124^I]1101 and [^124^I]1101-scFv8D3. Sagittal PET and autoradiography images of all Tg-ArcSwe and WT mice included in the study, acquired 72 h post injection of A) [^124^I]1101 or B) [^124^I]1101-scFv8D3, expressed as standardized uptake value (SUV, upper panel) or brain to blood ratio (middle panel). Note the different scales for the two antibody ligands. Ex vivo autoradiography (lower panel) from saline perfused, PET scanned mice

